# The involvement of neuroimmune cells in adipose innervation

**DOI:** 10.1101/787234

**Authors:** Magdalena Blaszkiewicz, Elizabeth Wood, Sigi Koizar, Jake Willows, Ryan Anderson, Yu-Hua Tseng, James Godwin, Kristy L. Townsend

## Abstract

**Background:** Innervation of adipose tissue is essential for the proper function of this critical metabolic organ. Numerous surgical and chemical denervation studies have demonstrated how maintenance of brain-adipose communication through both sympathetic efferent and sensory afferent nerves helps regulate adipocyte size, cell number, lipolysis, and ‘browning’ of white adipose tissue. Neurotrophic factors are growth factors that promote neuron survival, regeneration and plasticity, including neurite outgrowth and synapse formation. Peripheral blood immune cells have been shown to be a source of neurotrophic factors in humans and mice. Although a number of immune cells reside in the adipose stromal vascular fraction (SVF), it has remained unclear what roles they play in adipose innervation. We previously demonstrated that adipose immune cells secrete brain derived neurotrophic factor (BDNF).

**Methods:** We now show that deletion of this neurotrophic factor from the myeloid lineage led to a ‘genetic denervation’ of inguinal subcutaneous white adipose tissue (scWAT), thereby causing decreased energy expenditure, increased adipose mass, and a blunted UCP1 response to cold stimulation.

**Results:** We and others have previously shown that noradrenergic stimulation via cold exposure increases adipose innervation in the inguinal depot. Here we have identified a subset of myeloid cells that home to scWAT upon cold exposure and are Ly6C ^+^ CCR2 ^+^ Cx3CR1 ^+^ monocytes/macrophages that express noradrenergic receptors and BDNF.

**Conclusions:** We propose that these cold induced neuroimmune cells (CINCs) are key players in maintaining adipose innervation as well as promoting adipose nerve remodeling under noradrenergic stimulation, such as cold exposure.

## Introduction

In order for the central nervous system (CNS) to regulate functions of distal organs and tissues, peripheral nervous system (PNS) innervation needs to be maintained and properly coordinated. It has been demonstrated repeatedly that loss of innervation of the adipose organ (by surgical or chemical means) led to dysfunction of the tissue and disruption to whole body energy homeostasis. Denervation of brown adipose tissue (BAT) greatly impairs the energy expending process of adaptive thermogenesis (1–4), while denervation of white adipose tissue (WAT) results in fat mass accumulation via hyperplasia and impaired lipolysis (5–7). Furthermore, we have demonstrated that nerves in subcutaneous white adipose tissue (scWAT) undergo remodeling in response to environmental stimuli (cold, exercise), and exhibit signs of neuropathy in obesity/diabetes as well as with aging (8).

The maintenance of proper neural innervation is facilitated by neurotrophic factors (NFs) both in the CNS (9) and PNS (10). NFs are nerve growth factors that support nerve health, survival, and plasticity. Brain derived neurotrophic factor (BDNF) is one member of the neurotrophin family of NFs, which in mammals also includes nerve growth factor (NGF), Neurotrophin-3 (NT-3), and Neurotrophin-4/5 (NT-4/5). Neurotrophins signal predominantly through Trk receptors on nerves, through which they are endocytosed and, in peripheral nerves, transported in a retrograde manner to the nerve cell body, located in the ganglia. BDNF has been well studied for its role in hippocampal synaptic plasticity in the adult brain (11–13), as well as learning and exercise-related neurogenesis (14). BDNF is also an important modulator of energy balance through its actions in the central nervous system (15–17). Deletion of *Bdnf* in the ventromedial and dorsomedial regions of the hypothalamus resulted in an obesity phenotype due to hyperphagic behavior (16). Obesity is associated with lower serum levels of BDNF in humans (18, 19) while dietary restriction normalizes BDNF deficits in the brain in a mouse model of Huntington’s disease (20). Animal studies have shown that central and peripheral administration of BDNF reduced food intake and hyperglycemia and increased energy expenditure, via CNS mediated mechanisms (21–23). As review by Xu and Xie, genetic mutations in human BDNF and its receptor TrkB result in morbid early-onset obesity (24); furthermore, genome wide associated studies (GWAS) have identified single nucleotide polymorphisms (SNPs) in or near *BDNF* to be associated with increased body mass index (BMI) (24).

Despite these strong correlations between altered function of BDNF and its receptor TrkB with obesity, this growth factor has been predominately studied only in the CNS. A few studies have clearly shown that NFs, including BDNF, are present in adipose tissue (25–27), but the cellular source of BDNF in adipose had not been determined. In adipose specific (*FABP4-Cre*) knock-outs of BDNF or TrkB, only CNS effects were apparent (not surprising given off-target brain effects of this cre line) (28). Using instead an *adipoq*-cre (mature adipocyte-specific line) there was no obesity in a BDNF or a TrkB knock-out (KO), and the BDNF KO showed no difference in adipose BDNF levels (28). Together, these findings suggested that mature adipocytes are *<u>not</u>* the cellular source of BDNF (28), which we have confirmed in this study by demonstrating predominance of BDNF expression in the SVF. NFs are secreted by glial cells in the brain (29) and Schwann cells in peripheral tissues (30). However, other cell types, predominantly immune cells, are also known sources of NFs (31–35) but are less studied.

As part of the immune response to injury, immune cells are critical players in wound healing, regeneration, and remodeling of various tissues. They are an important component of the adipose organ where they modulate the inflammatory response, clear the tissue of apoptotic cells, and mediate adipose tissue remodeling during obesity through an influx of monocytes (undifferentiated macrophages), neutrophils, T cells, B cells and mast cells (36–41). Macrophages and their monocyte precursors are myeloid lineage immune cells and comprise the highest fraction of immune cells present in adipose tissue (42). They are highly heterogeneous cells that are polarized by environmental stimuli to evoke differential responses within a tissue, including secretion of cytokines.

In a simplistic paradigm, classically activated macrophages (M1) act in a pro-inflammatory manner, while alternatively activated macrophages (M2) produce an anti-inflammatory response and initiate tissue remodeling after injury. Both M1 and M2 cells retain phagocytic behavior. During obesity, M1 adipose tissue macrophages (ATMs) greatly increase in number without an influx of resolving M2 macrophages, thus contributing to a chronic state of tissue inflammation (43). Inflamed, insulin-resistant adipose tissue histology is characterized by macrophage crown-like structures surrounding hypertrophic, hypoxic and dying adipocytes. On the other hand, it has been suggested that cold-induced browning of adipose promotes an M2 phenotype in ATMs (43), possibly promoting tissue remodeling and potentially serving as a source of NFs in adipose tissue. Importantly, the immunology field has uncovered that M1 and M2 designations are an oversimplification, and in reality many immune cells express markers of both subtypes and may be inter-converting between the two polarities, but this has not yet been fully explored in adipose tissues.

Although myeloid cells, including monocytes and macrophages, from peripheral blood have been shown to store and release NFs (44), it is still unclear what role these immune cells play in peripheral innervation, and it remains unknown how and if adipose-resident and infiltrating immune cells are stimulated to release NFs that act locally. Microglia, the CNS resident myeloid cells that are most similar to macrophages, are an accepted source of BDNF in the brain (45) and increase secretion of BDNF in response to neuroinflammation (46). Microglial-derived BDNF in the CNS promotes hippocampal synaptic plasticity (45) and neurogenesis (47). We hypothesized that myeloid lineage cells may play a similar role in adipose tissue, and generated a myeloid specific BDNF knock-out mouse model by crossing *LysMCre^+/-^* and *BDNF^fl/fl^* mice. Here we report a role for myeloid derived BDNF in the specific maintenance of scWAT innervation, and identify a subpopulation of monocyte/macrophages (Ly6C^+^CCR2^+^Cx3CR1^+^) that infiltrate adipose tissue in response to cold stimulation and express BDNF.

## Materials & Methods

### Mice, Metabolic Phenotyping, and in vivo Analyses Animals

The following mouse strains were obtained from The Jackson Laboratory: C57BL/6J (Stock # 000664); LysMCre+/-(B6.129P2-Lyz2/J, Stock # 004781); BDNF^fl/fl^ (Bdnf^tm3Jae^/J, Stock # 004339); R26R-EYFP (B6.129X1-Gt(ROSA)26Sortm1(EYFP)Cos/J, Stock # 006148); Cx3CR1-EGFP (B6.129P2(Cg)-Cx3cr1tm1Litt/J, Stock # 005582). Animals with myeloid specific deletion of BDNF (*LysMCre^+/-^ ::BDNF^-/-^)* were generated in our facility by crossing mice heterozygous for the myeloid -specific cre transgene (LysMCre^+/-^) with mice homozygous for floxed BDNF. Reporter mice were generated by crossing LysMCre^+/-^ x BDNF^flfl^ x R26R-EYFP. Animals were housed 3-5 to a cage providing for socialization, in a monitored temperature and humidity-controlled environment with 12/12hr light/dark cycle. Cages were replaced weekly, Ad libitum access to food and water was maintained. For all studies animals were sacrificed using CO_2_ followed by cervical dislocation.

### Dietary Fat Interventions and Food Intake

Adult male *LysMCre^-/-^::BDNF^fl/fl^* (CON) *and LysMCre^+/-^ ::BDNF^-/-^* (KO) mice were fed a 45% HFD diet from Research Diets (New Brunswick, NJ) for up to 11 weeks. Mice were housed 2-3 per cage, at room temperature. Body weight was measured weekly. Food intake was measured daily for 7 days, then weekly until the end of the experiment. Adiposity was assessed at the end of the study by weighing intact adipose depots after surgical removal.

### CLAMS

Metabolic cage analyses were conducted in a Comprehensive Laboratory Animal Monitoring System (CLAMS; Columbus Instruments, Columbus, OH), for measurement of oxygen consumption (VO_2_) and carbon dioxide production (VCO_2_), from which both respiratory exchange ratio (RER), and energy expenditure (Heat) were calculated: *RER=VCO_2_/VO_2_*; *Energy expenditure (heat) = CV * VO_2_ cal/hr, where CV is the “caloric value” as given by CV = (3.815 + 1.232) * RER.* Animals were single housed in a bedding free cage, at room temperature on a 12hr light/dark cycle. Mice were acclimated for 24-48hrs, after which VO_2_, VCO_2_, RER, and Heat were measured every 15min for 3 days (72hrs). Waveform analysis of CLAMS data was performed by matching every 15min measurement across all three 24hr-cycles. Two-way repeated measures analysis of variance (RM, ANOVA) was performed for average VO_2_, VCO_2_, RER, and Heat per group. An uncorrected Fisher’s Least Significance Difference test was performed for each time point between dietary groups as a post-hoc test. Interaction P values are reported, which represent differences in 24hr data between groups, as well as multiple comparison results for differences which were only day/night phase specific.

### CL316,243 Injections

Adult (12-13 week old) male C57BL/6 mice received daily intraperitoneal (i.p.) injections of ADRβ3 agonist CL316,243 (Tocris Bioscience, Bristol, U.K.; Cat # 1499), at 1.0 mg/kg BW or an equivalent amount of sterile saline, for 10-14 days.

### Cold Exposure Experiments

All cold exposure was carried out in a diurnal incubator (Caron, Marietta, OH, USA) at 5°C, and a 12hr light/dark cycle. Animals were housed two to a cage and continuously cold exposed for 3 - 14 days.

### Glucose Tolerance Test (GTT)

Animals were fasted overnight for 16hr, after which they received an i.p. bolus injection of 1g/kg glucose. Blood glucose was measured using tail vein blood with a hand-held glucometer (OneTouch UltraMini, LifeScan, Milpitas, CA, Johnson & Johnson, New Brunswick, NJ), at time 0 and at intervals of 15 minutes, 30 minutes, and 60 minutes after injection.

### Western Blots

Protein lysates were prepared by homogenizing frozen whole adipose depots in RIPA buffer using a Bullet Blender (Next Advance, Averill Park, NY), followed by Bradford Assay, and preparation of equal-concentration lysates in Laemmli buffer. 60ug of protein was loaded per lane of a 10% polyacrylamide gel, and following gel running, proteins were transferred to PVDF membranes for antibody incubation. Primary antibodies used included: anti-PGP9.5 (Abcam, Cambridge, U.K. Cat. #10404 RRID:AB_297145 and #108986 RRID:AB_10891773) used at a 1:1000 and 1:500 dilutions respectively; anti-UCP1 (Abcam, Cambridge, U.K. Cat. #10983; RRID:AB_2241462) used at 1:1000 dilution, anti-TH (Millipore Cat. # AB152; RRID:AB_390204; Merck Millipore, Burlington, MA), anti-β-tubulin (Cell Signaling Technology, Danvers, MA, USA; Cat. # 2146; RRID:AB_2210545), were all used at 1:1000 dilution. Secondary antibody was anti-rabbit HRP (Cell Signaling Ct # 7074; RRID:AB_2099233), used at a 1:3000 dilution. Blots were visualized with enhanced chemiluminescence (ECL; Pierce) on a Syngene G:BOX. Protein expression of PGP9.5, TH, and UCP1 was normalized to either β-tubulin or β in and quantified in ImageJ.

### Collection of Adipose Secretions and BDNF ELISA

BAT depots were dissected, weighed and minced in a petri dish containing DMEM (high-glucose, serum-free). Minced tissue was transferred to a 15mL conical tube with 5mL DMEM (loosely capped to keep tissue oxygenated) and placed in a shaking water bath at 37°C. Secretions were collected at time 0, 1hr, 2hrs, and 3hrs (1mL collected from conical tube at each time point and replaced with 1mL fresh DMEM). Secretions were stored at −80°C until processing. For ELISA, protein secretions were concentrated using Amicon ® Ultra Centrifugal Filters, Ultracel ® -100K (Millipore, Burlington, MA USA; Cat. # UFC510096), per manufacturer’s instructions. Mouse BDNF PicoKine™ ELISA Kit (Boster Biological Technology, Pleasanton, CA, USA; Cat# EK0309) was used per manufacturer’s instruction to determine amount of BDNF present in adipose active secretions.

### Thyroid Hormone ELISA

Mouse sera were used to measure circulating levels of thyroxine (T4) and triiodothyronine (T3). Circulating concentrations were determined by Enzyme-Linked Immunosorbant Assays (ELISA) at Maine Medical Center Research Institute’s Core Facilities (Scarborough, ME).

### SVF Isolation

Bilateral whole inguinal adipose depots were quickly dissected and weighed. Tissue was minced in 37°C pre-warmed DMEM (high glucose, serum free) containing 2mg/mL Roche Collagenase A (Millipore-Sigma, St. Louis, MO; Cat# 10103586001) at a volume of 10mL/depot. Minced tissue with collagenase containing DMEM was placed in a 50mL conical tube and transferred to a shaking water bath (350/min rotation) at 37°C. Every 10min cells were dispersed by vortex and pipette mixing. Full dissociation was usually achieved within 2hrs, when adipocytes were clearly visible and all tissue was dissociated. Dissociated media was poured through 100um cell strainers, rinsed with DMEM and centrifuged at 500g for 10min to separate adipocytes from SVF. After centrifugation adipocytes were collected (found floating on top); remainder of DMEM was removed sparing the SVF pellet. SVF pellet was incubated with 500uL of RBC lysis buffer on ice for 2min, after which 2mL of DMEM with serum was added to stop lysis. Cells were centrifuged at 500g for 5min at 4°C, and either collected for RNA or resuspended in FACS/MACS buffer for cell sorting.

### Magnetic-activated Cell Sorting (MACS)

SVF from bilateral whole inguinal adipose depots was isolated as described above and resuspended in degassed buffer (1XPBS pH7.2, 0.5% BSA and 2mM EDTA). Single-cell suspensions were sorted on the MidiMACS Quadro magnetic-activated cell separator system (Miltenyi Biotec, Bergisch Gladbach, Germany) according to manufactures instructions. Briefly, cells were stained with primary PE-conjugated antibody, CD11b-PE (Cat #130-098-087); a 1:10 antibody dilution per 10^7^ cells was used. Cells were incubated for 10min at 4-8°C, washed, and centrifuged at 300g for 10min. Washed cell pellet was resuspended in a 1:10 dilution of anti-PE microbeads included in Anti-PE MultiSort Kit (Cat. #130-090-757). Following 15min incubation at 4-8°C, cells were washed and centrifuged as described above. Cells were resuspended in 500µL of buffer and passed through LS columns of the MACS separator. LS columns were prepped according to manufacturer’s suggestion. Cells were washed 3 times, collected cells (CD11b-) were collected. LS columns were removed from the MACS separator and flushed with 5mL of buffer to release the magnetically labeled cell fraction (CD11b+). MicroBeads were removed using MicroSort release reagent included in Anti-PE MultiSort Kit (Cat. #130-090-757). MicroBead free CD11b+ cell fraction was labeled for the secondary marker, Anti-F4/80-APC (Cat. #130-102-942), following the same procedure as described for the primary marker, except that Anti-APC MicroBeads (Cat. #130-090-855) were used.

### Fluorescence-activated Cell Sorting (FACS)

SVF from bilateral whole inguinal adipose depots was isolated as described above. For cell sorting, the following 5 marker panel was used with DAPI exclusion for viability: Anti-Mouse Ly6C_BV570 (HK1.4), Anti-Mouse CD11b PE (M1/70), Anti-Mouse CX3CR1 PercP5.5, SA011F11), Anti-Mouse CD45-PE-Cy7 (30-F11), Anti-Mouse CCR2 A700 (475301). Sorting was performed on a BD™ FACSAria II™ cell sorter with SVF gated on CD45 and CD11b; CD45+CD11b-represented the non-myeloid population; CD45^+^CD11b^+^ myeloid fraction was gated on Ly6C, followed by CCR2 and Cx3CR1.

### 20 Cell Surface Marker tSNE Analysis of Myeloid Diversity in Male and Female Mice

Adult (13-15 week old) male and female control mice (BDNF^fl/fl^) were cold exposed for 10 days in a diurnal incubator as described above. Following cold exposed SVF from bilateral whole inguinal adipose depots was isolated as described above. Single cell suspensions were treated with unlabeled FC receptor blocking antibody cocktail (CD16/CD32) in 50ul of FACS buffer (HBSS + 5mM EDTA + 2% FCS). Cells were then incubated in an “staining antibody cocktail” against 20 cell surface markers for 60min at 4°C in 100ul. Cells were then washed in 1mL of FACS buffer and centrifuged at 400g for 7min, 2 times. Cells were then resuspended and analyzed on a five-laser 30-parameter FACSymphony A5 cytometer (BD Biocsiences, San Jose, USA) using DAPI exclusion for cell viability. For compensation of fluorescence spectral overlap, UltraComp eBeads (eBioscience, Inc.) were used following the manufacturer’s protocols. FCS 3.0 files generated by flow cytometry were initially processed using FlowJo Software (Tree Star, Ashland, USA) for automated compensation. Standard manual hierarchical gating was performed to remove debris, cell doublets and (DAPI+) dead cells from analysis before gating on CD45+ leukocyte populations. In preparation for performing the 20 marker tSNE leukocyte population analysis, the CD45+ population from each sample was down-sampled to 3000 events to normalize cellular input between samples. Using FlowJo, a concatemer of all samples was performed. An unbiased T-Distributed Stochastic Neighbor Embedding (tSNE) plugin algorithm was then run using defaults with 22 parameters, on the whole sample pool to obtain a multi-sample population reference map. Gating of each sample and experimental group was performed to generate tSNE maps for each condition. Differential cell clusters were gated and 20 marker histograms plots were used to predict cluster identities.

Antibodies used for adipose flow cytometric panel shown here: Anti-Mouse CD45-BUV395 (30-F11), Anti-Mouse B220_BUV496 (RA3-6B2), Anti-Mouse Ly-6G BUV563 (Clone 1A8), Anti-Mouse CD19 BUV661 (Clone 1D3), Anti-Mouse CD11b BUV737 (M1/70), Anti-Mouse NKp46_BV421 (29A1.4), Anti-Mouse CD62L BV510 (MEL-14), Anti-Mouse Ly6C_BV570 (HK1.4), Anti-Mouse CD3 BV605 (17A2), Anti-Mouse Mrc1_BV650 (C068C2), Anti-Mouse MHCII -I-A/I-E -BV711 (M5/114.15.2), Anti-Mouse NK1.1 BV785 (PK136), Anti-Mouse CD11c_FITC (HL3), Anti-Mouse CD80 PE (16-10A1), Anti-Mouse CD115 CF594 (AFS98), Anti-Mouse CX3CR1 PercP5.5 (SA011F11), Anti-Mouse CD64 PE-Cy7 (X54-5/7.1), Anti-Mouse CD14 APC (Sa2-8), Anti-Mouse CCR2 A700 (475301), Anti-Mouse F480 APC/Cy7 (BM8).

### Gene Expression (qPCR)

RNA was isolated from whole tissue depots using Trizol reagent, and total RNA extracted using a Zymo (Irvine, CA) kit. RNA yield was determined on a Nanodrop; cDNA was synthesized using a High Capacity Synthesis Kit (Applied Biosystems, Foster City, CA). Real-time quantitative (q)PCR was performed with SYBR Green (Bio-Rad, Hercules, CA) on a CFX96 instrument (Bio-Rad, Hercules, CA).

### Histology

#### Adipose

Immunofluorescent staining of 10% buffered formalin fixed, paraffin-embedded, 7um sections of adipose tissues was performed for detection of UCP1 (RRID:AB_2241462, 1:500, Abcam, Cambridge, UK. Cat. #10983). Alexa 488 (2.5µg.mL, Molecular Probes, Eugene, OR, USA, Cat. # A11070) was used as secondary antibody. Typogen Black staining was used to quench tissue autofluorescence (prior to antibody incubation) and also provided visualization of cell size, browning (multilocularity) and crown-like structures. Stained sections were mounted using Millipore mounting fluid (Burlington, MA USA; Cat. # 5013) and 1 1/2 coverslips, and imaged on a Nikon Eclipse E400 epifluorescent microscope equipped with Nikon DS-fi2 camera. Wholemount adipose staining was performed as previously described (8, 48) and imaged by widefield and confocal microscopy. Widefield imaging was performed on a Nikon E400 epifluorescence microscope with Hamamatsu ORCA-Flash4.0 V2 Digital CMOS monochrome camera (Hamamatsu Photonics K.K., C11440-22CU). Objectives used: Nikon CFI Plan Apochromat Lambda 10X (NA 0.45, WD 4.00mm, dry) and Nikon CFI Plan Fluor 40X (NA 0.75, WD 0.66mm, dry.) Images were processed using Nikon NIS-Elements software. Images captured on Hamamatsu ORCA-Flash4.0 V2 were pseudo colored. For confocal imaging a Leica TCS SP8 DLS microscope with HyD detectors was used. Fluorophores were excited with a white light laser tuned specifically for the excitation and emission spectra of EGFP. Objectives used: HC PL APO CS2 20X (NA 0.75, WD 0.62mm, dry) and HC PL APO CS2 63X (NA 1.40, WD 0.14mm, oil). Images were processed with LASX software and were either pseudo colored or left monochromatic. Primary antibodies included: PGP9.5 conjugated to Alexa 647 (1:200, Abcam, Cambridge, U.K. Cat. #196173).

#### Neuromuscular Junction Immunofluorescence, Imaging, and Analysis

Soleus and medial gastrocnemius muscles were removed and fixed in a 2% PFA at 4°C for 2 hours. Tissues were rinsed with 1XPBS and incubated in blocking buffer (1XPBS/2.5%BSA/0.5-1%Triton) at 4°C for at least 24 hours and up to 7 days. Following blocking muscles were teased, tendons and fat was removed, and tissue was flattened by being placed between two tightly-bound glass slides for at least 30 minutes at 4°C. Tissues were transferred to fresh blocking buffer at 4°C for at least 12 hours. Immunostaining of innervation with primary antibodies was performed overnight at 4°C, followed by 1XPBS washes on a rotating platform at 4°C replacing PBS every 1hr for a total of 4-6hrs. Tissues were incubated with secondary fluorescent antibodies in similar fashion as primary antibodies. Primary antibodies included: neurofilament-M (2H3, RRID:AB_531793, 1:500) and synaptic vesicles (SV2, RRID:AB_2315387, 1:250) from Developmental Studies Hybridoma Bank, (University of Iowa, USA). Secondary antibodies included: Alexa Fluor 488 at 1:500 (A21121) and alpha-bungarotoxin (BTX)- conjugated to Alexa Fluor 594 at 1:1000 (B13423) from Molecular Probes (Eugene, OR, USA). Tissues were mounted on microscope slides using Millipore mounting fluid (Burlington, MA USA; Cat. # 5013) and 1 1/2 coverslips then sealed and allowed to set overnight. Stained sections were imaged on a Nikon Eclipse E400 epifluorescent microscope equipped with Nikon DS-fi2 camera. Brightness, contrast, and sharpness were adjusted in Microsoft PowerPoint. Up to 100 NMJs were counted for each tissue, statistics were conducted in GraphPad PRISM software (La Jolla, CA, USA) using the multiple t-tests (one-per row) function.

### Statistical Analysis

For all animal experiments, mice were randomized to treatment groups to ensure no difference in starting body weight. All plots represent mean +/-SEM. Statistical calculations were carried out in Excel or GraphPad Prism software (La Jolla, CA, USA), utilizing ANOVA, Linear Regression, or Student’s T-test as indications of significance (specified in Figure legends). Gene and protein expression data were normalized to a housekeeper and analyzed by either ANOVA or by Student’s t-test, two-tailed, using Welch’s correction when variance was unequal. Error bars are SEMs. For all figures, *p < 0.05, **p < 0.01, ***p < 0.001, ****p < 0.0001.

### Ethical Statement

All procedures and handling of animals were performed in accordance with the University of Maine’s Institutional Animal Care and Use Committee (IACUC), to comply with the guidelines of the PHS Policy on Humane Care and Use of Laboratory Animals, and Guide for the Care and Use of Laboratory Animals. This study was approved by the University of Maine’s IACUC, under protocol A2017-09-04.

## Results

### BDNF is expressed primarily in adipose SVF

We previously demonstrated that BDNF secretion increases from scWAT in response to noradrenergic stimulation, after administration of the β-3 adrenergic receptor (ADRβ3) agonist CL316,243 (8). We then endeavored to determine which adipose compartment and cell type was the source of local BDNF secretion in scWAT. Adipose tissue is a heterogeneous organ and although adipocytes are the main cell type, numerous other cell types are contained within the stromal vascular fraction (SVF) of adipose tissue. Adipose SVF consists predominantly of hematopoietic lineage cells including adipose tissue macrophages (ATMs (41, 42)), but also contains numerous other immune cell types, preadipocytes, vascular endothelial cells, and pericytes. To determine the compartmental source of adipose BDNF, adult male C57BL/6 mice were cold exposed and whole SVF was isolated from mature adipocytes of inguinal scWAT by collagenase dispersion. Gene expression analysis revealed that *Bdnf* was almost exclusively expressed in the SVF, while other common growth factors (nerve growth factor; NGF, and vascular endothelial growth factor, VEGFa) showed no difference of expression between SVF and mature adipocyte fractions (Fig. 1A).

**Figure 1.**
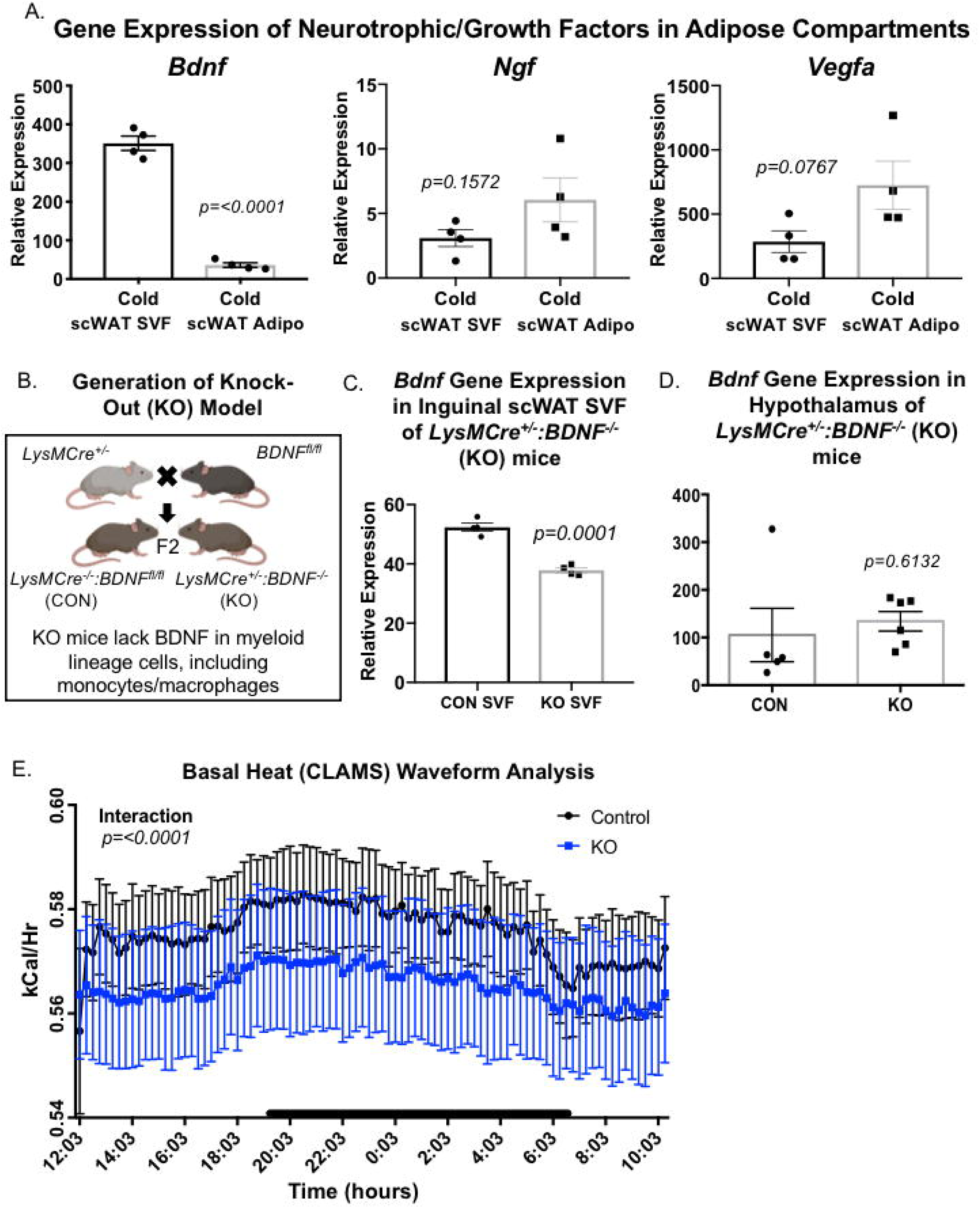
BDNF is expressed in scWAT SVF fraction; *LysMCre^+/-^::BDNF^-/-^* (KO) mice have lower energy expenditure. Adult male CB57/BL6 mice were cold exposed at 5°C and SVF was isolated from mature adipocytes of the inguinal subcutaneous white adipose (scWAT) depot. Differences in gene expression between adipose compartments of neurotrophic factors, *Bdnf, Ngf,* and *Vegfa,* is shown (A). Data analyzed by Student’s t-test, two-tailed, N=4. Illustration of knock-out model generation (B). BDNF gene expression in adipose SVF of *LysMCre^-/-^::BDNF^fl/fl^* (CON) versus *LysMCre^+/-^::BDNF^-/-^* (KO) mice (C). Data analyzed by Student’s t-test, two-tailed, N=4 per group. Gene expression of *Bdnf* in hypothalamus of *LysMCre^-/-^::BDNF^fl/fl^* (CON) versus *LysMCre^+/-^ ::BDNF^-/-^* (KO) mice (D). Data analyzed by Student’s t-test, two-tailed, N=5 CON; N=6 KO. Adult (8-12 week old) male CON and KO mice were assessed in metabolic cages (CLAMS). KO mice displayed lower energy expenditure represented as heat calculated from measures of VO_2_ and VCO_2_ over the whole 24hr (E). Waveform analysis of metabolic cage measurements taken at 15min increments for 48 hrs. Time of day is indicated on the x-axis, animals were maintained on a 12 hr light/dark cycle (black bars indicate dark cycle). For all error bars are SEMs.

Given the known expression of BDNF in immune cells, we sought to determine the contribution of myeloid-lineage BDNF to adipose innervation by creating a knock-out (KO) mouse model using Cre-Lox technology. *LysMCre^+/-^* mice were bred to *BDNF^fl/fl^* mice to generate *LysMCre^+/-^::BDNF^-/-^* (KO) animals, which lacked BDNF in myeloid lineage cells (Fig. 1B). (For the genotyping strategy, see Suppl. Fig. S1A.) Compared to littermate controls, KO animals exhibited a significant, although not complete, decrease of *Bdnf* in adipose SVF as measured by gene expression (Fig. 1C). Likely additional SVF cell types also contribute to tissue BDNF levels. Since myeloid lineage cells are also expressed in the brain, and BDNF has been shown to play a role in energy balance via CNS control of satiety, we also investigated whether our KO model affected BDNF expression in the hypothalamus. Gene expression of *Bdnf* in the hypothalamus did not differ between KO animals and their littermate controls (Fig. 1D). Physiological assessment using metabolic cages performed on KO animals and their littermate controls in the basal state revealed that KO mice had significantly lower energy expenditure (Fig. 1E) despite showing no difference in body weight or adiposity (Suppl. Fig. S1B).

### KO mice have a blunted response to cold stimulation

Since a decrease in energy expenditure could be indicative of impaired sympathetic drive, we stimulated sympathetic nerve activity via cold exposure. When adult (22-23 week old) male mice were cold challenged at 5°C for 4 days, KO and control animals maintained similar body weight (Fig. 2A, left panel). However, KO animals had a trend for increased adiposity (Fig. 2A, right panel). As cold exposure stimulates catecholamine-induced lipolysis mediated through sympathetic nerves, we next investigated innervation of the inguinal scWAT depot following a 7-day cold challenge. Protein expression of the pan-neuronal marker PGP9.5 was markedly reduced in inguinal scWAT of male KO mice compared to littermate controls (Fig. 2B, left panel). Protein expression of tyrosine hydroxylase (TH), a marker of sympathetic innervation and activation, was also drastically reduced in inguinal scWAT of KO animals compared to controls (Fig. 2B, right panel). Gene expression of synaptic/innervation markers (*Psd95, Sox10, Synapsin I, Synapsin II, Synaptopysisn)* in inguinal scWAT also showed a coordinated trend to be decreased in KO animals (Suppl. Fig S1C), further indicating perturbations in scWAT innervation of KO animals. Together, these data indicated a ‘genetic denervation’ in this model.

**Figure 2.**
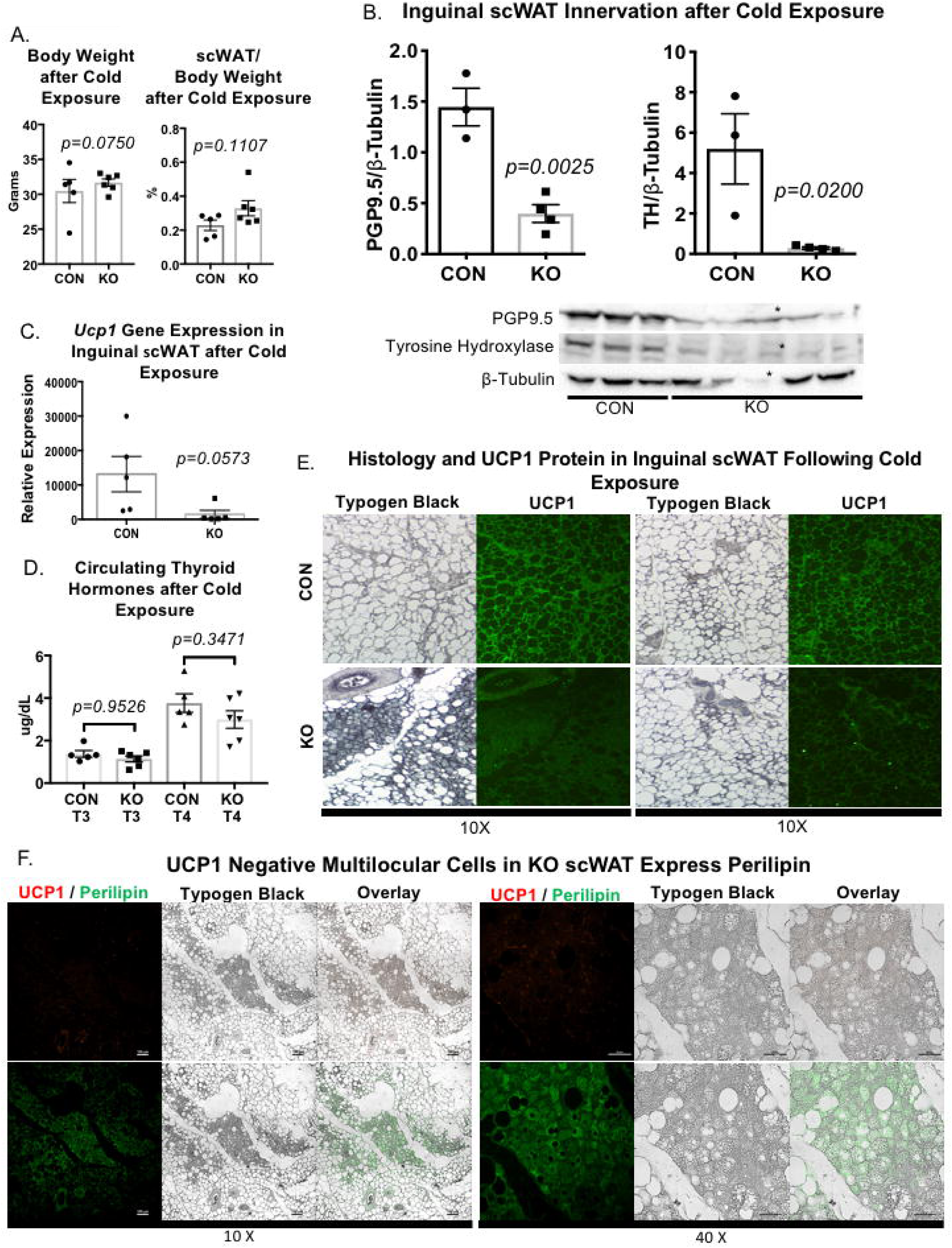
*LysMCre^+/-^::BDNF^-/-^* (KO) have increased adiposity and impaired response to cold due to genetic denervation of scWAT. Adult (22-23 week old) male *LysMCre^-/-^::BDNF^fl/fl^* (CON) and *LysMCre^+/-^ ::BDNF^-/-^* (KO) mice were cold exposed at 5°C for 4 days; body weight and adiposity were compared between CON and KO groups (A). Body and tissue weight data were analyzed by two-tailed Student’s T-Test, N=5 CON, N=6 KO. Protein expression of PGP9.5 and tyrosine hydroxylase (TH) in inguinal scWAT was measured by western blotting from adult (12-25 week old) 7-day cold (5°C) exposed WT/CON and KO male animals (B). β-tubulin was used as a loading control for normalization. Data were analyzed by two-tailed Student’s T-Test, N=3 WT/CON, N=4 KO, * denotes data that was removed from analysis due to lack of expression of loading control. Gene expression of *Ucp1* was measured in adult (12-25 week old) 7-day cold (5°C) exposed WT/CON and KO males (C). Data were analyzed by two-tailed Student’s T-Test, N=5 WT/CON, N=5 KO. Circulating thyroid hormones, thyroxine (T4) and triiodothyronine (T3), were measured by ELISA from serum of adult (22-23 week old) 4-day cold (5°C) exposed CON and KO male mice (D). Data were analyzed by One-way ANOVA, with Tukey’s multiple comparisons test, N=5 CON, N=6 KO. Immunofluorescent staining for UCP1 was performed on inguinal scWAT sections of adult (22-23 week old) male CON and KO mice following 4-day cold (5°C) exposure (E). Immunofluorescent staining for UCP1 and Perilipin was performed on inguinal scWAT sections of adult (22-23 week old) male KO mice following 4-day cold (5°C) exposure (F). Typogen Black, used to quench lipid autofluorescence, provided staining cell morphology which was visualized under brightfield microscopy. Overlay is immunofluorescence over brightfield of the same area (F). Images were acquired with a 10X or 40X objective and are representative of N=5 CON, N=8 KO (E) and N=4 KO (F). For all error bars are SEMs.

Interestingly, gene expression of lipolytic markers *Atgl* and *Hsl* showed an increased trend in scWAT of KO animals (Suppl. Fig S1C). Loss of scWAT innervation potentially results in decreased lipolysis due to lack of SNS mediated release of norepinephrine. However, due to the critical role of lipolysis in facilitating thermoregulation during cold challenge, we suspect that lipolysis in KO animals is achieved by alternative, nerve-independent, pathways. For example, glucocorticoid-mediate lipolysis has been attributed to increased β-adrenergic responsiveness via direct signaling or indirectly by inducing secretion of angiopoietin-like 4 (Angptl4) (49, 50).

Cold exposure induces expression of uncoupling protein 1 (UCP1) as required for adaptive thermogenesis, due to its ability to uncouple the mitochondrial respiratory chain resulting in a proton leak and heat production. UCP1 is therefore a unique marker of BAT and cold-induced ‘browning’ in scWAT. Gene expression of *Ucp1* in inguinal scWAT of 7-day cold exposed mice was reduced in KO animals compared to littermate controls (Fig. 2C). Thyroid hormone potentiates sympathetic nervous system (SNS) activation of thermogenesis in BAT, but is produced via a distinct neuronal pathway from adipose sympathetic drive. We observed no changes in circulating thyroid hormones between KO and control (CON) animals (Fig. 2D). Histological assessment of inguinal scWAT revealed what appeared like increased multilocularity suggestive of browning in KO animals (Fig. 2E, Typogen Black staining). However, there was a striking lack of UCP1 expression in these areas in KO tissues (Fig. 2E). Typogen Black (used here to reduce tissue autofluorescence) may also mark immune cells, thus there is potential for increased immune cell infiltration in the KO tissues as well. However, perilipin (a lipid droplet associated protein) staining of scWAT from KO animals clearly showed that the multilocular cells lacking UCP1 expression contained high amounts of lipid droplets (Fig. 2F, UCP1 immunofluorescence in red, perilipin immunofluorescence in green on serial sections). Preadipocytes maintain multilocular lipid droplets prior to differentiation, and considering that denervation of WAT increases hyperplasia, the observed multilocularity could more likely be areas of increased preadipocytes. In total, these findings demonstrate reduced thermogenic potential in denervated scWAT.

### Genetic denervation of scWAT in LysMCre^+/-^::BDNF^-/-^ (KO) mice is depot specific

We next sought to determine whether genetic denervation of scWAT in KO animals was restricted to this adipose tissue or extended to other organs. BDNF is a known myokine, and muscle is an energy expending tissue. We assessed innervation of fast twitch (gastrocnemius) and slow twitch (soleus) muscle in CON and KO, by investigating occupancy of neuromuscular junctions (NMJs) at basal conditions in adult male mice. Immunostaining of the presynaptic nerve and vesicles (neurofilament and SV2 markers, respectively) with postsynaptic acetylcholine receptors was performed to allow visualization of NMJ (Suppl. Fig. S2A, left panel). Following counts of occupied, partially occupied, and unoccupied NMJs it was determined that there was no evidence of neurodegeneration in the NMJ of KO animals (Suppl. Fig. S2A, right panel). In the same animals we also assessed axon numbers of spinal (L5 ventral root), motor, and sensory nerves through cross-section imaging (Suppl. Fig. S2B). A lower axon count could reflect neuronal death in these larger PNS nerves, but no difference was observed between CON and KO animals.

Myeloid cells are also present in BAT tissue. We therefore wanted to evaluate whether a lack of BDNF in BAT myeloid cells would have an effect on this tissue’s function. BAT of 7-day cold exposed 12-25 week old male CON and KO mice was evaluated. Protein expression of neuronal markers TH and PGP9.5 did not differ between KO and control mice (Suppl. Fig. S3A), indicating that BDNF may not have an important neurotrophic role in BAT. Consistent with that, histological assessment of BAT revealed no difference in cellular morphology nor UCP1 expression (Suppl. Fig. S3B). Indeed, when adult male C57BL/6 mice were cold exposed or treated with the pharmacological ADRβ3 agonist, CL316,243, no difference in BDNF secretion from BAT was observed when compared to basal conditions (Suppl. Fig. S3C). Together, these data supported scWAT depot specificity of our genetic denervation model, and BAT neuronal function may simply be due to increased sympathetic nerve activity and not changes in neural plasticity and neurite outgrowth in the tissue. Alternatively, a different neurotrophic factor, like NGF, may be more important for BAT than WAT.

### HFD feeding exacerbates fat mass accumulation in LysMCre^+/-^::BDNF^-/-^ (KO) Mice

Loss of sympathetic innervation to inguinal scWAT has been shown to increase depot mass (51), however, in our genetic model of scWAT denervation, no difference in adiposity was observed under basal conditions (Suppl. Fig. S1B) despite the observed decrease in energy expenditure (Fig. 1E). We next metabolically challenged CON and KO mice with a 45% high fat diet (HFD). Adult (25 week old) male CON and KO mice were placed on a HFD for 3-11 weeks, to assess adipose integrity and energy balance.

At 3 weeks of HFD feeding animals were characterized in metabolic cages. HFD resulted in only a slight decrease in energy expenditure in KO mice compared to littermate controls (Fig. 3A) as analyzed via waveform analysis over multiple 24hr periods. However, KO mice showed a higher respiratory exchange ratio (RER) than CON animals during the light cycle, indicative of preferential metabolism of carbohydrates over lipids for fuel (Fig. 3B). These data fits with studies demonstrating that adipose nerves are important for lipolysis (52) and that denervation would shift fuel preference to carbohydrates. These physiological differences between CON and KO animals were observed despite no difference in food intake or change in body weight (Fig. 3C-D) at early time points. After 6 weeks of HFD, KO mice displayed altered glucose control compared to CON animals (Fig. 3E), perhaps due to a shift in fuel utilization. By 11 weeks of HFD feeding, KO animals displayed greater adiposity than littermate controls (Fig. 3F), potentially due to the reduced innervation and loss of neural control of certain metabolic processes.

**Figure 3.**
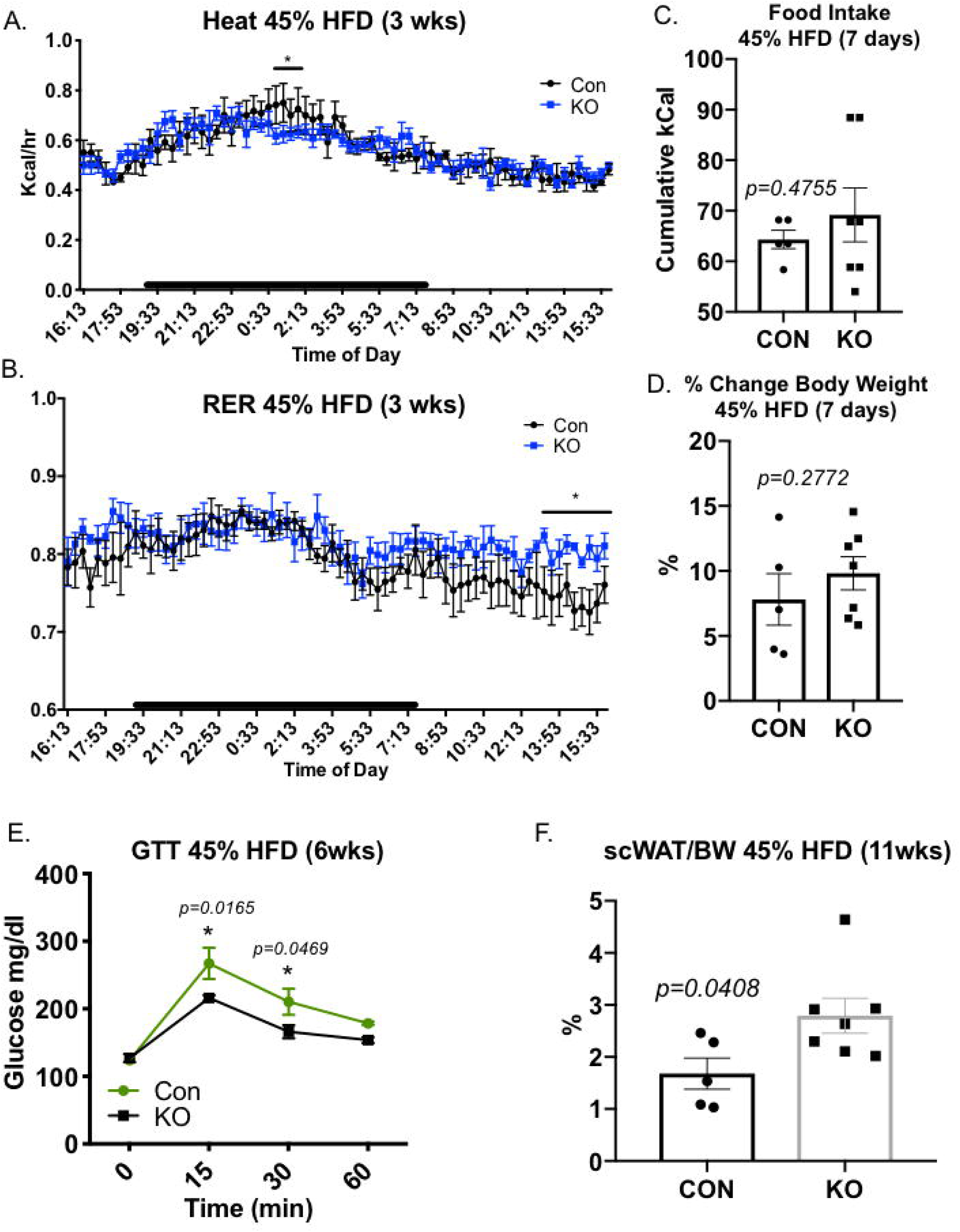
*LysMCre^+/-^ ::BDNF^-/-^* (KO) showed accelerated fat accumulation on a 45% HFD. Adult (25 week old) male *LysMCre^-/-^::BDNF^fl/fl^* (CON) and *LysMCre^+/-^ ::BDNF^-/-^* (KO) were challenged with a 45% HFD for 3 weeks before undergoing physiological assessment in metabolic cages (A-B). Energy expenditure as measured by heat was lower for KO versus CON only for a short period during the dark cycle (A). Respiratory exchange as a ratio (RER) between the two groups, indicated greater use of carbohydrates for fuel by KO animals during the light cycle (B). Data presented as waveform analysis of measurements taken at 15min increments for 3 days. Time of day is indicated on the x-axis, and animals were maintained on a 12 hr light/dark cycle (black bars indicate dark cycle). Data analyzed by two-way repeated measures ANOVA with Fisher’s LSD test; N=4 per group. Adult (25 week old) male CON and KO animals were placed on a 45% HFD, daily food intake (represented as cumulative food intake) (C) was measured for the first week of HFD feeding. Percent change in body weight (D) was measured for the first 7 days of HFD feeding. Data were analyzed by two-tailed Student’s T-Test, N=5 CON, N=7 KO. Glucose tolerance testing was performed at 6 weeks of HFD feeding (E). Data were analyzed by Two-way ANOVA, with Tukey’s multiple comparisons test, N=5 CON, N=5 KO. Adiposity was measured for CON and KO animals after 11 weeks of HFD feeding as a percentage of scWAT over body weight (F). Data were analyzed by two-tailed Student’s T-Test, N=5 CON, N=7 KO. For all error bars are SEMs. *p < 0.05, **p < 0.01, ***p < 0.001, ****p < 0.0001.

### Cold-induced neuroimmune cells (CINCs) are recruited to scWAT and express BDNF

After demonstrating the significance of myeloid derived BDNF to scWAT innervation, we sought to determine which myeloid cells were the source of tissue BDNF. Given their multifaceted role in adipose tissue, being a source of BDNF in the brain (microglia), and the phenotype observed in a myeloid-lineage KO, we hypothesized that monocytes/macrophages were the leading source of BDNF in scWAT. Since previous studies indicated that BDNF is increased in scWAT with noradrenergic stimulation (8), CD11b^+^ F4/80^+^ macrophages were isolated from SVF of inguinal scWAT of room temperature and 5-day cold exposed C57BL/6 adult (12 week old) male mice. *Bdnf* gene expression did not differ between room temperature and cold exposed CD11b^+^ F4/80^+^ macrophages (Fig. 4A). F4/80+ is considered a pan-macrophage marker, as such we considered it too broad to reveal phenotypic changes in the spectrum of macrophage populations in scWAT. Furthermore, F4/80+ is not the ideal marker of monocytes (macrophage precursors), which could be infiltrating the tissue in response to cold exposure. Based on these findings we applied a different approach to determining which myeloid cells are the source of scWAT BDNF. Adult (12 week old) female control animals were maintained at room temperature or cold exposed for 10 days. Inguinal scWAT SVF was isolated and flow cytometrically analyzed using a custom antibody cocktail against immune cells. Surprisingly, cold exposure did not have an effect on either M1 or M2 ATMs (Fig 4B). Instead, the greatest increase was in Ly6C^+^CCR2^+^ monocytes. Both Ly6C^+^ and CCR2^+^ are markers of inflammatory monocyte migration (53). Following cold exposure, both Ly6C^+^CCR2^+^Cx3CR1^-^ and Ly6C^+^CCR2^+^Cx3CR1^+^ populations increased in inguinal scWAT of female mice, however, only the Ly6C^+^CCR2^+^Cx3CR1^+^ population showed a strong trend for increase in males (Fig. 4C). In a separate cohort of adult male and female C57BL/6 mice, unbiased assessment using t-distributed Stochastic Neighbor Embedding (tSNE) revealed subpopulations of immune cells changing in propensity in scWAT with cold, and between male and female mice (cold and room temperature). Interestingly, the Ly6C^+^CCR2^+^Cx3CR1^+^ population, which we call cold-induced neuroimmune cells (CINCs) again increased in both male and female scWAT with cold (Fig. 4D).

**Figure 4.**
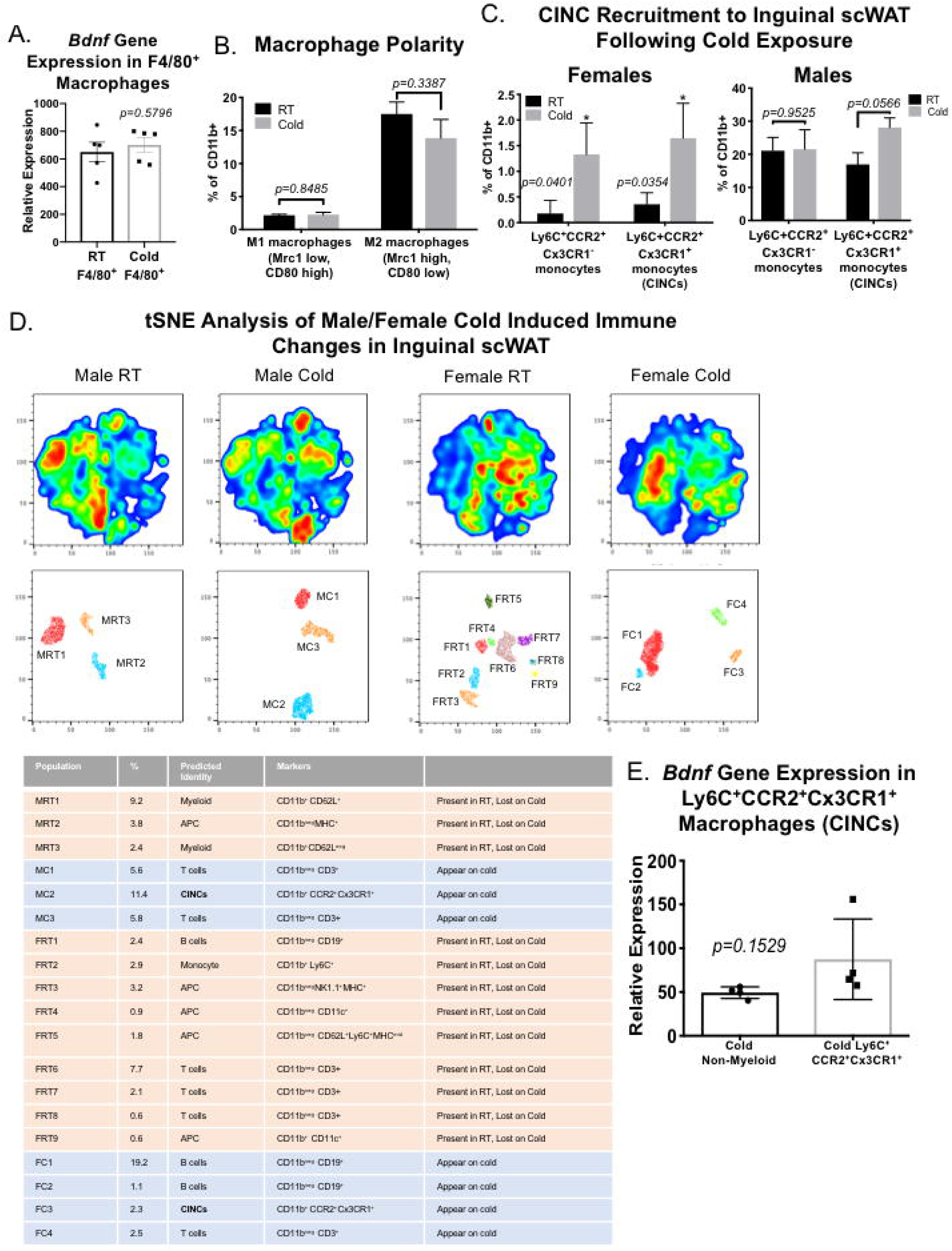
Cold induced neuroimmune cells (CINCs) home in to inguinal scWAT and express *Bdnf*. Adult (12 week old) male C57BL/6 were either maintained at room temperature (RT) or cold exposed (5°C) for 5 days, ATMs from inguinal scWAT depots were isolated using magnetic-activated cell sorting (MACS) by positive selection of CD11b^+^ followed by F4/80^+^ cells. *Bdnf* gene expression in doubly labeled CD11b^+^ F4/80^+^ macrophages was compared between RT and cold exposed animals (A). Data were analyzed by two-tailed Student’s T-Test, N=4 per group. Adult (12 week old) female control animals were either maintained at room temperature (RT) or cold exposed (5°C) for 10 days, SVF from bilateral inguinal scWAT was isolated and FACS sorted using a 20 cell surface marker panel for myeloid lineage immune cells (B-C). Changes in M1/M2 polarity (B) and Ly6C^+^CCR2^+^Cx3CR1^-^ and Ly6C^+^CCR2^+^Cx3CR1^+^ macrophage precursors/monocytes (C, left panel) were measured between RT and cold exposed animals. Adult (12-13 week old) male C57BL/6 were either maintained at room temperature (RT) or cold exposed (5°C) for 14 days; SVF from bilateral inguinal scWAT was isolated and FACS sorted; Ly6C^+^CCR2^+^Cx3CR1^-^ and Ly6C^+^CCR2^+^Cx3CR1^+^ cells were compared between RT and cold exposed animals (C, right panel). Data were analyzed by two-tailed Student’s T-Test, N=3-4 per group for both sexes. Adult male (M) and female (F) control animals were either maintained at room temperature (RT) or cold exposed (5°C) for 10 days, SVF from bilateral inguinal scWAT was isolated and FACS sorted using a 20 cell surface marker panel for myeloid lineage immune cells (N=5 per group). t-Distributed Stochastic Neighbor Embedding (tSNE) analysis was performed to identify myeloid lineage cell population changes in response to cold exposure; Ly6C^+^CCR2^+^Cx3CR1^+^ were identified for both sexes as adipose cold induced neuroimmune cells (CINCs) (D). *Bdnf* gene expression measured in Ly6C^+^CCR2^+^Cx3CR1^+^ cells indicated that these infiltrating cells express *Bdnf* (E). Data analyzed by two-tailed Student’s T-Test, N=4 per group. For all error bars are SEMs. *p < 0.05, **p < 0.01, ***p < 0.001, ****p < 0.0001.

To confirm that Ly6C^+^CCR2^+^Cx3CR1^+^ monocytes/macrophages were the source of BDNF, we FACS sorted out Ly6C^+^CCR2^+^Cx3CR1^+^ cells from inguinal scWAT SVF of 14 day cold exposed adult (12-13 week old) male C57BL/6 mice. We measured *Bdnf* gene expression in cold induced Ly6C^+^CCR2^+^Cx3CR1^+^ cells and found that they showed a trend for increased expression of *Bdnf* compared to cold exposed non-myeloid cells (Fig. 4E). We believe the previously observed increase in BDNF in adipose SVF ((8) and Fig. 1A) was due to infiltration of CINCs homing to scWAT upon cold exposure, and not an increase in per-cell BDNF levels.

*Adr*β*3* gene expression in Ly6C^+^CCR2^+^Cx3CR1^+^ cells confirmed the presence of norepinephrine (NE) receptor on these cells, indicating the potential to be responsive to SNS stimulation (Suppl. Fig. S4A), a likely mode for promoting BDNF release to the tissue after cold exposure. Taken together, these data indicated that Ly6C^+^CCR2^+^Cx3CR1^+^ cells are cold-induced neuroimmune cells (CINCs) that increase in number in scWAT after cold, have the potential to be stimulated by sympathetic nerves, and express BDNF.

### Adipose lymph node and adipose lymphatics may play a role in recruitment of CINCs

CINCs express the transient markers Ly6C^+^ and CCR2^+^, as well as Cx3CR1^+^, indicating that they are recruited to adipose tissue following cold stimulation as opposed to being tissue resident immune cells. Immune cells are generally recruited to tissue through vasculature upon chemoattract release from target tissue. To further investigate dynamics of CINC recruitment to adipose tissue we utilized a Cx3CR1-EGFP reporter mouse. Whole mount imagining of axillary adipose tissue clearly demonstrated that Cx3CR1^+^ macrophages were present at the surface of and within the adipose lymph node (LN; Fig. 5A-B). 3D reconstruction of adipose LN imaging with depth coding allowed us to visualize Cx3CR1^+^ macrophages present on the capsule as well as within the cortex of the LN (Fig. 5B, middle and right panel). Furthermore, Cx3CR1^+^ macrophages were enriched in adipose lymphatics (Fig. 5C left panel, yellow arrow) compared to blood vasculature (Fig. 5C left panel, red arrow). We took advantage of innate autofluorescence to visualize vasculature in these samples, and although blood and lymph vasculature are somewhat morphologically similar, the bulbous sacs found at capillary terminals are a unique feature of lymphatic vasculature only (Fig. C middle and right panels, bulbous sacs outlined in white). These lymphatics were the vessels that predominantly contained Cx3CR1^+^ cells, and not the blood vasculature.

**Figure 5.**
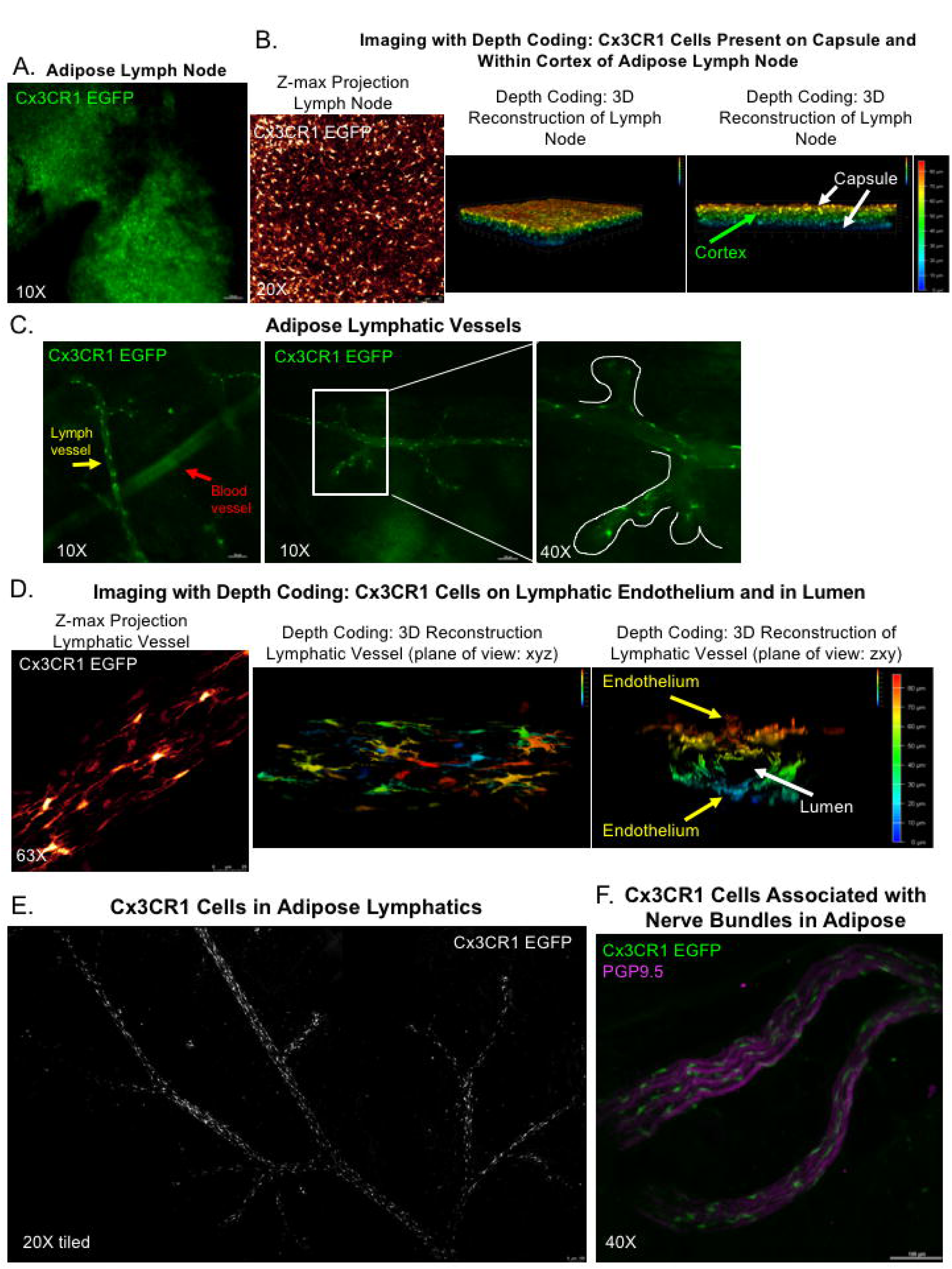
Cx3CR1 Cells in Adipose Transit via Lymphatics. Wholemount imaging of female (A-E) and male (F) axillary scWAT from Cx3CR1-EGFP reporters mice. Cx3CR1+ cells (green) in lymph node captured by widefield microscopy at 10X (A). Cx3XR1+ cells were shown to occupy the entire depth of the lymph node, captured by confocal microscopy at 20X (B), represented as a z-maximum projection (Glow LUT) (B, left) and 3D reconstruction (depth coded) at two angles (B, middle and right.) Cx3CR1+ cells (green) line lymphatic vasculature (morphologically distinguished from blood vasculature by the bulbous sacs on initial lymphatics, yellow arrow and outlined in white) but are absent in blood vasculature (red arrow) (C). Captured by widefield microscopy at 10X and 40X. Cx3CR1+ cells (Glow LUT) imaged by confocal microscopy at 63X were shown to reside on lymph vessel endothelium and were present within the lumen (D). Z-maximum projection of lymph vessel (D, left) and 3D reconstruction (depth coded) at two angles: looking down z-axis (D, middle) and looking down the x-axis (D, right) with vessel lumen identified by arrow. Tiled z-maz projection captured on confocal at 20X demonstrates that Cx3CR1+ cells (white) are found throughout the lymphatic network in scWAT (E). Cx3CR1+ cells (green) reside around nerve bundles marked by the pan-neuronal marker PGP9.5 (magenta) as captured by widefield microscopy at 40X (F).

Confocal imaging of Cx3CR1^+^ macrophages on lymphatic endothelium clearly demonstrated that these cells have a distinct morphology (Fig. 5D left panel). The cell bodies are elongated with cytoplastic extensions creating a spindeloid shape associated with alternatively activated macrophages (54). 3D reconstruction with depth coding following confocal imaging of adipose lymph vessels revealed that Cx3CR1^+^ macrophages did not merely interact with lymph endothelial cells but were present within the lumen of the vessels (Fig. 5D, middle and right panels). Cx3CR1^+^ macrophages were present in greater numbers around and within the lymphatics of adipose (Fig. 5E), and as previously reported were also associated with adipose nerves (Fig. 5F) (55). Taken together, these data indicated that Cx3CR1+ macrophages are transported by the lymphatic vasculature to scWAT.

## Discussion

Here we present evidence of the necessity for myeloid-derived BDNF in maintaining inguinal scWAT innervation. Loss of BDNF from LysM^+^ myeloid cells resulted in a decrease in total innervation of the inguinal adipose tissue as measured by protein levels of the pan-neuronal marker PGP9.5, and a near complete denervation of sympathetic nerves (as measured by TH protein expression). We attribute this ‘genetic denervation’ as a cause for the observed decrease in energy expenditure exhibited under basal conditions, impaired thermogenic potential evidenced by lack of UCP1 induction and increased WAT mass during cold stimulation, and worsened response to a high-fat, high-sugar diet. These phenotypes fit with the known roles of adipose nerves in regulating proper adipogenesis, lipolysis, thermogenesis and overall metabolic health in adipose tissues.

Diet-induced obesity results in chromic low grade adipose inflammation due to infiltration of pro-inflammatory immune cells to the tissue, and ATM content increase from 10-15% (in lean mice) to ∼50% with obesity (36, 56). In addition, phenotypic switching of CD4+ T cells and recruitment of T cell and B cells precedes macrophage infiltration, and with obesity macrophage polarity skews to a more inflammatory phenotype.

We initially hypothesized that anti-inflammatory (M2) ATMs were the source of scWAT BDNF. However, we observed no changes in M1 or M2 populations in scWAT after cold exposure. These M1 and M2 designations are an oversimplification of the diversity of macrophage populations and represent diverse subsets of cells with unique molecular signatures. Instead of changes in M1 or M2, we saw an increase in Ly6C^+^CCR2^+^Cx3CR1^+^ monocytes in scWAT of both male and female cold-exposed mice. We have named these cells CINCs since they are recruited to scWAT by cold and express BDNF. Surprisingly, it would appear that CINCs are a pro-inflammatory cell type as Ly6C^+^ and CCR2^+^ are markers of inflammatory monocytes. Although pro-inflammatory cell infiltration to adipose tissue is usually a harbinger of metabolic dysfunction, an acute inflammatory response is necessary in wound healing and tissue remodeling. Only chronic, ongoing inflammation would be detrimental and causative of insulin resistance.

On the other hand the, the Cx3CR1^+^ macrophages we identified in adipose lymphatics exhibited features (such as elongated cell bodies) that are morphologically attributed to M2 or anti-inflammatory macrophages (54) further underscoring the incredible diversity of macrophage characterization in adipose. By utilizing an unbiased 20 antibody labeling approach to assess adipose SVF, we have discovered subsets of myeloid cells that respond to these particular environmental stimuli, providing further granularity to the myriad of immune cells types active in adipose.

Recently, a CX3CR1^+^ population of macrophages had been described in association with sympathetic nerves in adipose tissue (55). These sympathetic nerve associated macrophages (SAMs) were found to regulate catecholamine levels on adipose tissue by phagocytosing and degrading norepinephrine (NE) (55). Whether SAMs are the same cells as our CINCs remains uncertain. Similar to CINCs, SAMs *<u>do</u>* exhibit pro-inflammatory markers, and a distinct morphology similar to the Cx3CR1^+^ macrophages we identified in the adipose lymphatics. SAMs home to WAT in the obese state, while conversely, we see CINCs home to WAT with cold/noradrenergic-stimulation. Given that these two populations home to WAT in different conditions, this could indicate that the two immune cell types may have opposing roles, including potential for phagocytosing nerve debris for SAMs and stimulating nerve regeneration for CINCs. Fitting with this view, SAMs sequester and degrade NE, while CINCs express BDNF.

On the other hand, CxCR1^+^ macrophages have been shown to play diverse and even opposing roles in the intestines (57), a phenotypic plasticity which may also be present in adipose tissue. One thing that is clear is the M1/M2 paradigm of ATM classification is an oversimplification of the functionally distinct populations of macrophages active in adipose tissue. Further markers beyond Cx3CR1^+^ are necessary to fully understand and phenotype the rich variety of macrophages in adipose. Compounding the difficulty in clearly delineating macrophage populations is that macrophage activation is a dynamic process. Phenotypic switching appears to be sequential, responding to microenvironment stimuli (58) and may be dependent on spatiotemporal differences in tissue resident immune cell subtypes (59).

Since CINCs home into adipose tissue, we used a Cx3CR1-EGFP reporter mouse to visualize how this happens. Immune cells are recruited to tissues via vasculature, however, we found it surprising that in adipose it appeared that lymphatic and not blood vasculature plays a more important recruitment role for CINCs. Strikingly, the majority of Cx3CR1^+^ cells were contained within the lymph node and lymphatic vessels of adipose tissue. Because lymph flows in a unidirectional manner, the lymphatic system is often overlooked as a source of bidirectional crosstalk. In fact, lymphatic cells are not all destined for unidirectional trafficking. Leukocytes can also exit lymphatic vasculature and enter surrounding tissues (60).

We have previously reported increased innervation around LNs in adipose following cold exposure (8), while others have shown that browning commonly occurs around LNs (61). Furthermore, all lymph nodes in the body are surrounded by adipose tissue, which is activated during an immune response (62). Together these data underscore the potential crosstalk between nerves, adipocytes, and leukocytes in scWAT. Regardless of whether or not CINCs are recruited to or merely drained by lymphatic vessels, the importance of the lymphatic system in transporting adipose immune cells is again underscored by our current findings.

We postulate that CINCs contribute to nerve remodeling under noradrenergic stimulation, leading to BDNF release (Suppl. Fig. S4B). Cao, *et al.* have recently suggested that sympathetic nerve plastically is dependent on cold-induced, adipose-derived NGF (63). Although they provide evidence that NGF is involved to some degree in promoting cold induced sympathetic nerve density and browning of scWAT, the cellular source of NGF was ambiguous (our data indicate it is produced more by mature adipocytes than SVF cells, and thus may be promoting innervation of the adipocytes and not the SVF). After determining that NGF gene expression increased in the first couple of days of cold exposure, Cao, *et al.* used an NGF neutralizing antibody to prevent NGF activity and observed decreased sympathetic nerve density in response to cold. Their approach in preventing TrkA receptor function in adipose nerves and showing decreased browning and sympathetic nerve density lends credence to the notion that NGF plays a role in adipose innervation, similar to the role we observe for BDNF. Taken together it is clear that locally-derived nerve growth factors are important for maintaining proper brain-adipose communication and adipose innervation for metabolic control. Considering the variety of NFs and the multitude of cell types in adipose tissue it would not be surprising if both NGF and BDNF play a role in adipose innervation through different mechanisms. Promoting innervation of a specific axonal subtype to target a specific cell in the tissue may in fact be accomplished by coordinated secretion of a given growth factor by the target cell type; whereby mature adipocytes secrete NGF to ‘wire up’ adipocytes to TrkA+ axons, whereas SVF immune cells secrete BDNF to ‘wire up’ immune cells to TrkB+ axons. The role of BDNF in peripheral nerve health is often overlooked in lieu of the overwhelming focus on central BDNF action. However, BDNF is known to play an important role in peripheral nerve regeneration, as reviewed by *McGregor & English* (64). Following peripheral nerve injury local sources of secreted BDNF include not only dorsal root ganglia neurons but support cells, including Schwann cells (64). Cx3CR1+ macrophages have also been shown to play a crucial role in peripheral nerve regeneration by regulating Schwann cell dynamics in mice and humans (65). These studies align with the data presented here and offer a new avenue of research in neuroimmune regulation of peripheral nerve plasticity.

## CONCLUSIONS

In conclusion, we have demonstrated the importance of CINC immune cells for maintaining adipose innervation, particularly after cold exposure, which promotes CINC movement to adipose (likely via adipose lymphatic vasculature). CINCs express and likely secrete BDNF, thus promoting axonal survival and metabolic function in the tissue. CINCs also express adrenergic receptor, making them a likely target for cold-stimulated NE release in order to promote BDNF secretion to scWAT, once they have homed to the tissue. Given these findings, it is clear that neuroimmune crosstalk in scWAT is important for tissue function and metabolic control.

## Availability of data and materials

Data available upon request.

## Acknowledgements

The authors wish to thank the following individuals from University of Maine: Brenda Kennedy-Wade and Prof. James Weber for assistance in the animal facility; Dawna Beane for paraffin embedding; Callie Greco, Cameron Fudge, and Sarah Berez for animal care. The authors also wish to thank Drs. Gregory Cox and Robert Burgess from Jackson Laboratory for sharing their NMJ assessment methods; Dr. Maribel Rios for expert advice; and Nicholas Cutter and Emma Bragdon for help with spinal, motor, and sensory nerve assessments.

## List of abbreviations

ADRβ3: β-3 adrenergic receptor
ATMs: Adipose tissue macrophages
BAT: Brown adipose tissue
BDNF: Brain derived neurotrophic factor
BMI: Body mass index
CINCs: Cold induced neuroimmune cells
CNS: Central nervous system
CON: Control
FACS: Fluorescence-activated cell sorting
GWAS: Genome wide associated studies
HFD: High fat diet
KO: Knock-out
LN: Lymph node
MACS: Magnetic-activated cell sorting
NE: Norepinephrine
NFs: neurotrophic factors
NGF: nerve growth factor
NMJs: Neuromuscular junctions
NT-3: Neurotrophin-3
NT-4/5: Neurotrophin-4/5
PGP9.5: Protein gene product 9.5
RER: Respiratory exchange ratio
SAMs: Sympathetic nerve associated macrophages
scWAT: Subcutaneous white adipose tissue
SiLN: Subiliac lymph node
SNPs: Single nucleotide polymorphisms
SNS: Sympathetic nervous system
SVF: Stromal vascular fraction
TH: Tyrosine hydroxylase
tSNE: T-distributed Stochastic Neighbor Embedding
UCP1: Uncoupling protein 1
VEGFa: Vascular endothelial growth factor a
WAT: White adipose tissue

## Funding

For this project, the Townsend Lab was funded by start-up funds from the Office of the Vice President for Research at University of Maine, a Rising Tide Seed Grant from University of Maine, a Junior Faculty Award from the American Diabetes Association (1-14-JF-55), and an NIH NIDDK 1R01DK114320-01A1. The Godwin lab is supported by a NIH COBRE award (P20GM104318).

## Author Contributions

MB designed experiments, analyzed data, and wrote the manuscript. EW and SK collected and analyzed histology data. SK and RA collected and analyzed data related to brown adipose tissue. JG conducted the flow cytometric analysis on adipose immune cells. YHT oversaw initial pilot experiments. KLT designed experiments, analyzed data, conceived of and oversaw the project, and wrote the manuscript. KLT is the guarantor of this work and, as such, had full access to all the data in the study and takes responsibility of the data and the accuracy of the data analysis.

## Declarations

The authors do not have any conflicts of interest to disclose.

- **Ethical Approval and Consent to participate:** All animal studies were IACUC approved at University of Maine
- **Consent for publication:** All authors have consented for publication.

## Supplemental Information

Supplemental Figure S1

Supplemental Figure S2

Supplemental Figure S3

Supplemental Figure S4

Supplemental Figure Legends

## Notes

### Competing Interest Statement

The authors have declared no competing interest.

