## Supplemental Text for "The involvement of neuroimmune cells in adipose innervation"

**Supplemental Figure S1.** **Basal body and adipose of** ***LysMCre^+/-^::BDNF^-/-^* (KO) animals and gene expression of inguinal scWAT following cold exposure.**  Representative genotyping gel for determining *LysMCre^+/-^::BDNF^-/-^* (KO) versus *LysMCre^-/-^::BDNF^fl/fl^* (CON) (A). For the BDNF foxed allele a band is expected at 487bp, wild type band is expected at 437bp; therefore fl/fl denotes homozygous floxed, fl/+ denotes heterozygous floxed and +/+ denotes wild type. Cre allele is detected at 337bp, the band at 438bp serves as a PCR control expected in all DNA samples; -/- denotes lack of cre, +/- indicates presence of cre allowing recombination to occur. Animals A-B are designated as *LysMCre^-/-^::BDNF^fl/fl^* (CON), animals C-F are designated as *LysMCre^+/-^::BDNF^-/-^* (KO). Adult (29 week old) male *LysMCre^-/-^::BDNF^fl/fl^* (CON) and *LysMCre^+/-^::BDNF^-/-^* (KO) were assessed under basal condition for body weight and subcutaneous adiposity (B). Data analyzed by two-tailed Student’s T-Test, N=4 CON, N=6 KO. Gene expression of synaptic markers (*Psd95, Sox10, Synapsin I, Synapsin II, Synaptophysin*) and lipolytic markers (*Atgl, Hsl*) in inguinal scWAT following 4-day cold (5°C) exposure in adult (22-23 week old) male mice (C). Data analyzed by Student’s t-test, two-tailed, N=4 CON; N=5 KO.

**Supplemental Figure S2. Further phenotyping of *LysMCre^+/-^::BDNF^-/-^* (KO) animals.** Immunostaining of the presynaptic nerve and vesicles (neurofilament and SV2, in green) and postsynaptic acetylcholine receptors (in red) was performed on fast twitch (medial gastrocnemius) and slow twitch (soleus) muscles to allow visualization and assessment of the neuromuscular junction (NMJ) (A). Data analyzed by two-tailed Student’s T-Test, and representative of male 23-25 week old CON and KO mice, N=3 per group. For the same group, axon numbers of spinal, motor, and sensory nerves were counted through cross-section electron microscopy imaging (D). Error bars are SEMs

**Supplemental Figure S3. Assessment of BAT in *LysMCre^+/-^::BDNF^-/-^* (KO) animals.** Adult (12-25 week old) male *LysMCre^-/-^::BDNF^fl/fl^* (CON) and *LysMCre^+/-^::BDNF^-/-^* (KO) animals were cold (5°C) exposed in diurnal chambers for 5-7 days (A-B). Protein expression of tyrosine hydroxylase (TH) and PGP9.5 in BAT was compared between CON and KO mice (A). Data analyzed by two-tailed Student’s T-Test, N=6 CON, N=8 KO. Immunostaining for UCP1 (green) was performed on paraffin embedded cut sections of BAT from CON and KO animals (B). Typogen Black staining shows cell morphology, images taken with 40X objective for total magnification of 400X. Data representative of N=6 CON, N=8 KO. Adult (12-13 weeks old) male C57BL/6J mice were maintained at room temperature (RT) or cold exposed (5°C) for 10 days, or received either daily i.p. injections of ADRβ3 agonist CL316,243 (at 1.0 mg/kg BW), or vehicle for 14 days. *Ex vivo* secretions (collected at 1hr and 2hr) from BAT explants were measured for BDNF by ELISA, and not analyzed due to low N of control animals, N=2 RT/VEH, N=4 Cold/CL (C).

**Supplemental Figure S4. *Adrβ3* gene expression in CINCs.** Adult (12-13 week old) male C57BL/6 were either maintained at room temperature (RT) or cold exposed (5°C) for 14 days; SVF from bilateral inguinal scWAT was isolated and FACS sorted to isolate CINCs. *Adrβ3* gene expression was measured in RT and cold exposed Ly6C^+^CCR2^+^Cx3CR1^+^ cells (A). Data analyzed by two-tailed Student’s T-Test, N=4 per group. For all error bars are SEMs. Model illustrating the involvement of neuroimmune cells in adipose innervation (B).
