## Supplementary figures and images for "The involvement of neuroimmune cells in adipose innervation"

### Supplemental Figure 1

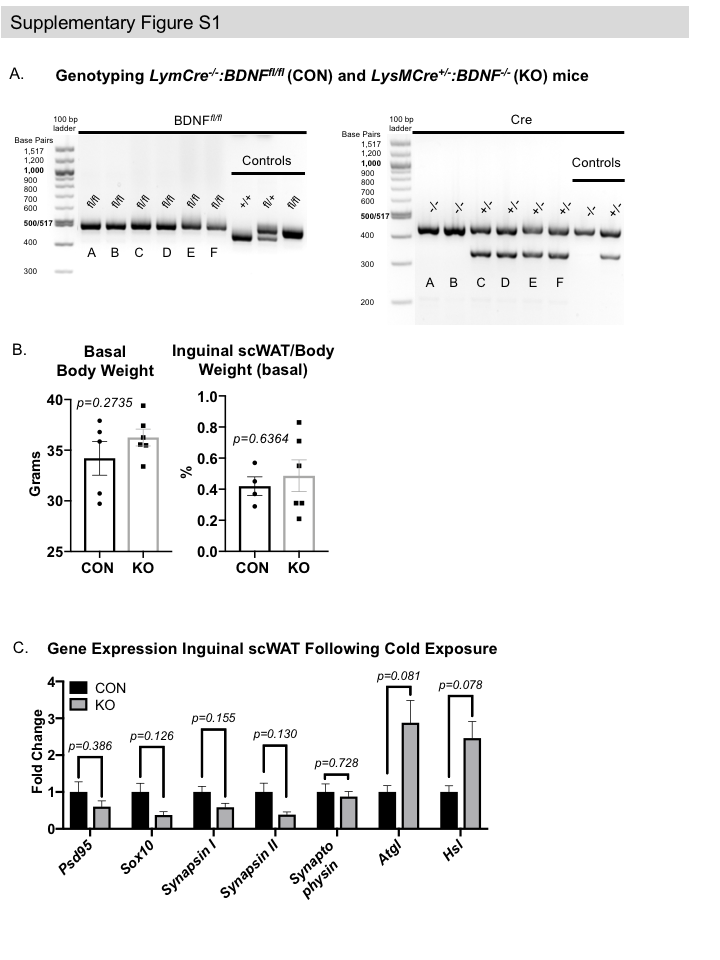

### Supplemental Figure 2

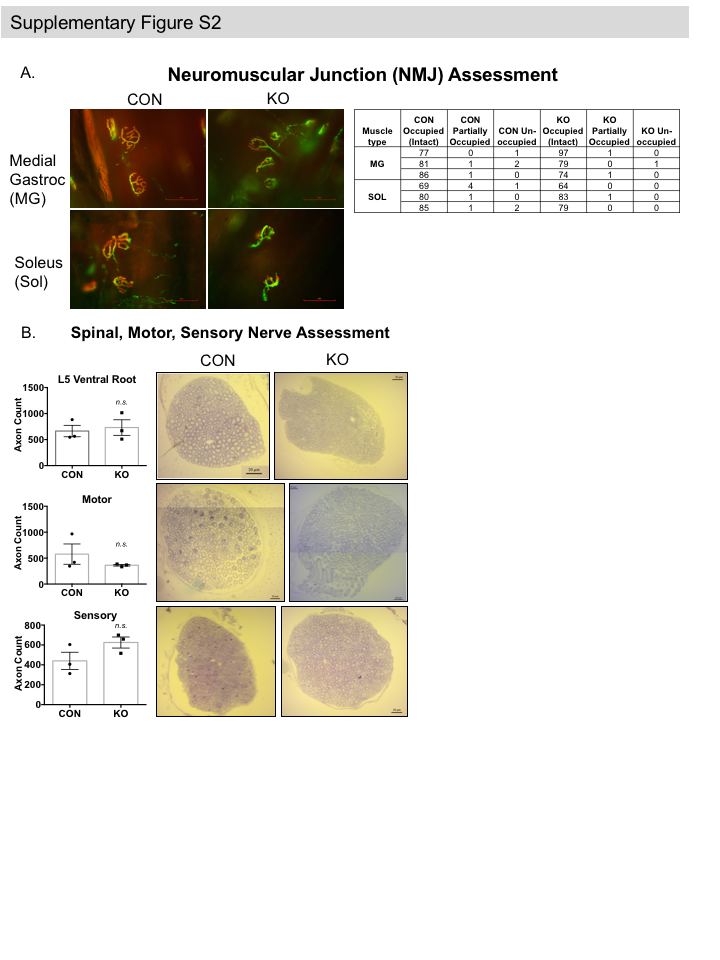

### Supplemental Figure 3

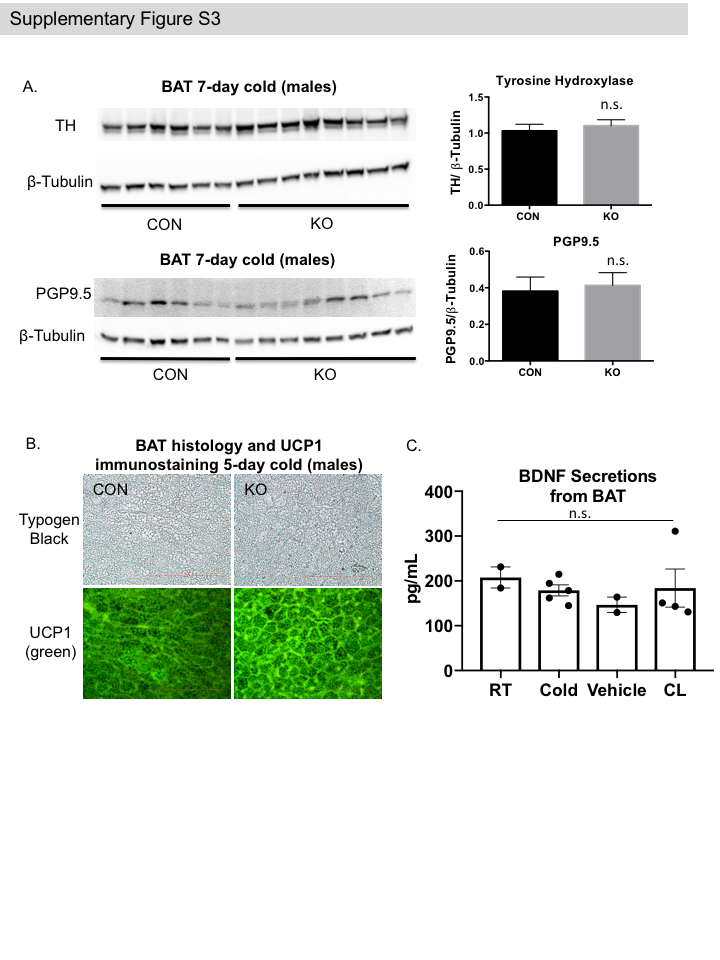

### Supplemental Figure 4

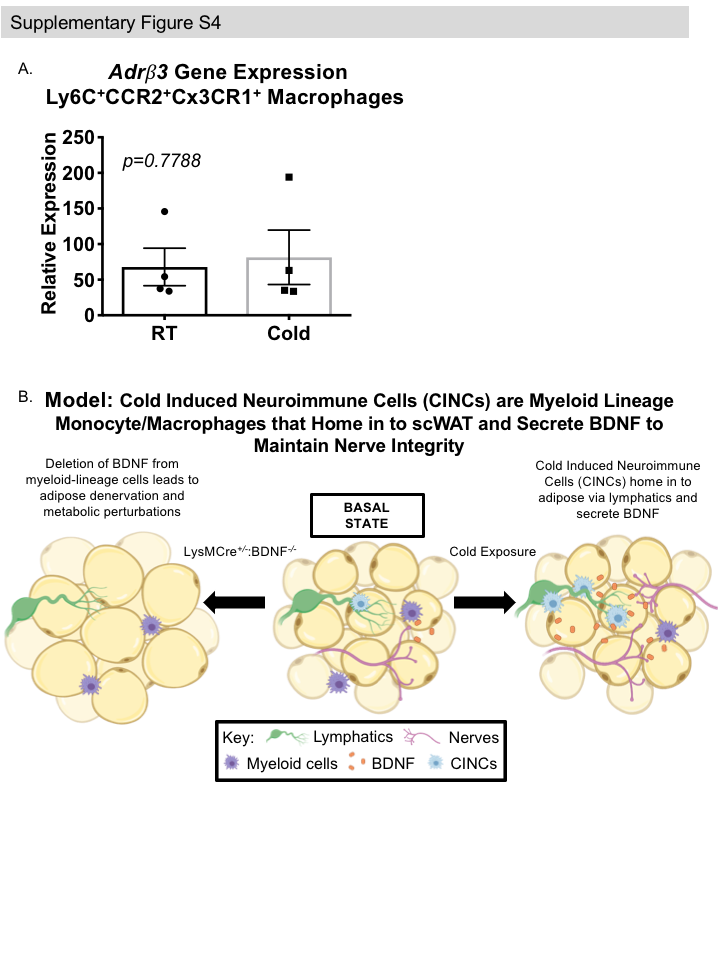
